# Cell competition reprograms adherent-suspension plasticity in disseminated tumor cells

**DOI:** 10.1101/2024.11.25.625320

**Authors:** Hyunbin D. Huh, Yujin Sub, Jee Hung Kim, Hannah Lee, Jongwook Oh, Dong Ki Lee, Jong Hwa Byun, Woo Yang Pyun, Sehyung Pak, Junjeong Choi, Tadayoshi Hashimoto, Takayuki Yoshino, Joon Jeong, Heon Yung Gee, Hyun Woo Park

## Abstract

Cell competition within the primary tumor drives tumor growth by promoting the uncontrolled proliferation of winner cells and eliminating loser cells by sensing cell fitness. However, the mechanism of how cell competition confers the loser cells with the metastatic potential of circulating tumor cells (CTCs) to transit between dissemination and colonization remains elusive. Here we found cell competition gives rise to less fit but viable cells that transformed into CTCs via adherent-to-suspension transition (AST) mechanisms. The induction of hematopoietic transcription factors hijacked by solid tumor cells rendered CTCs competent to reprogram their anchorage dependency and disseminate into the bloodstream, while subsequent suppression of these factors was critical to regain adhesion and colonize metastatic lesions. Disrupting the oscillatory dynamics of AST factors blocked the adherent-suspension plasticity (ASP) of breast cancer CTCs and suppressed lung metastasis. Furthermore, multiregional single-cell transcriptomic analyses of matched primary tumors, CTCs, and metastatic lesions from *de novo* metastatic breast cancer patients demonstrate the critical role of ASP in metastasis.

**Significance:** We demonstrate cell competition-mediated displacement of loser cells manifests dynamic oscillation of AST factors that confer anchorage plasticity to circulating tumor cells critical for their dissemination and colonization in metastasis. These findings highlight the potential of targeting AST factors to develop effective anti-metastatic therapies.

## Introduction

Cell competition in multicellular organisms plays a crucial role in maintaining tissue homeostasis by eliminating less-fit cells and allowing competitive cells to thrive. This fitness-driven selection process operates through mechanical competition that depends on cell-cell interactions or molecular competition driven by secreted factors by neighboring cells. In primary tumors, cell competition is increasingly recognized as a driving force of evolution, facilitating clonal expansion of aggressive cells while displacing weaker cells from the tumor niche. Intriguingly, recent evidence suggests that a rare population of displaced, less-fit cells may not simply be passively eliminated but rather actively reprogrammed to contribute to metastasis and recurrence.

Metastasis, the primary cause of cancer death, involves the dissemination of primary tumor cells into the bloodstream as circulating tumor cells (CTCs), their survival within the circulation, and eventual colonization of distant organs. The metastatic cascade, therefore, could be perceived as a consequence of cancer cells constantly transitioning between an adherent state and a suspension state. Based on these observations, we and others recently discovered a biological phenomenon referred to as adherent-to-suspension transition (AST), a mechanism orchestrated by the induction of specific hematopoietic transcription factors, collectively termed AST factors, that reprograms the anchorage dependency of solid tumor cells to facilitate the dissemination and release of CTCs via downregulation of cell-matrix adhesion pathways and induction of anoikis resistance by mitigating oxidative stress. However, whether cell competition during cancer progression confers dynamic plasticity of AST-mediated hematopoietic mimicry to disseminated tumor cells that endows successful completion of the metastatic cascade remains unknown.

Here, to investigate the role of adherent-suspension plasticity (ASP) in CTCs during the dissemination and colonization processes, we recapitulated cell competition *in vitro* through dissemination assay, which enabled a rare population of cancer cells to undergo spontaneous detachment and reattachment from a highly confluent and unfavorable cell culture condition that mimics the tumor microenvironment. Surprisingly, these *in vitro* CTC-like cells showed dynamic oscillation of AST factor expression, which emerged during detachment and subsequently suppressed in the course of reattachment. We further show that *in vitro* CTC-like cells reflect similar characteristics to *in vivo* AST-positive CTCs derived from an orthotopic model of breast cancer metastasis.

The establishment of advanced patient cohorts that consist of matched primary tumors, CTCs, and metastatic lesions from combined tissue and liquid biopsy specimens provides invaluable insights to reveal undescribed mechanisms, targets, and theories of metastatic plasticity. Thus, moving forward from current multi-omics analyses^1–5^, we newly enrolled untreated *de novo* metastatic breast cancer patients, collected matched tissue and liquid biopsy specimens and analyzed single-cell level transcriptomes of paired primary tumors, CTCs, and metastatic lesions. It was necessary to compare the transcriptome of CTCs with their matched solid tumor counterparts harvested from the same patient to monitor the dynamic plasticity of AST factors and related gene signatures within the metastatic cascade from dissemination to colonization. Thus, through the development of a cell competition assay and analyzing single-cell-level transcriptomes of matched specimens from *de novo* metastatic breast cancer patients, our study reveals how anchorage plasticity drives the metastatic cascade via AST factor-mediated ASP in disseminated CTCs.

## Results

### Cell competition reprograms loser cells with AST-mediated anchorage plasticity

Metastasis could be conceived as a longitudinal sequence of cancer cells constantly transitioning between their adherent and suspension states. Solid tumors shed CTCs through a fitness-based cell competition mechanism within the overcrowded environment of the primary tumor. Here, we asked whether the rare population of displaced, less-fit cells are actively reprogrammed to confer metastatic traits of CTCs rather than simply being passively eliminated.

To examine the mechanism underlying the displacement of loser cells from adherent parental cells, we induced cell competition by challenging various subtypes of breast cancer cell lines, including MDA-MB-231, MCF7, BT549, and Hs578T cells, to high cell density stress conditions. Notably, many breast cancer cell lines subjected to overcrowded cell competition exhibited cell-cell contact inhibition or cell death. In MDA-MB-231 cells, however, cell competition induced cell rounding and spontaneous detachment of CTC-like cancer cells that resemble the rare population of disseminated CTCs *in vivo* (**Fig. 1A-B**). We hypothesized that the ability of a cell line to produce CTC-like cells through cell competition would likely reflect its metastatic capacity *in vivo*. To test this, we injected each breast cancer cell line into the fat pad of mice and examined their metastatic efficiency to disseminate and colonize the lung (**Fig. 1C**). As expected, despite showing similar primary tumor growth, the cell lines that failed to generate CTC-like cells via cell competition assay showed significantly lower metastatic potential and prolonged survival compared to MDA-MB-231 (**Fig. 1D**). These observations are in line with earlier reports that classified MDA-MB-231 as one of the most metastatic breast cancer cell lines in mouse models (**Fig. S1A**) ^6^. Thus, these results suggest that the metastatic potential of breast cancer cells correlates with the ability to produce spontaneously suspended CTC-like cells via cell competition assay.

**Figure 1.**
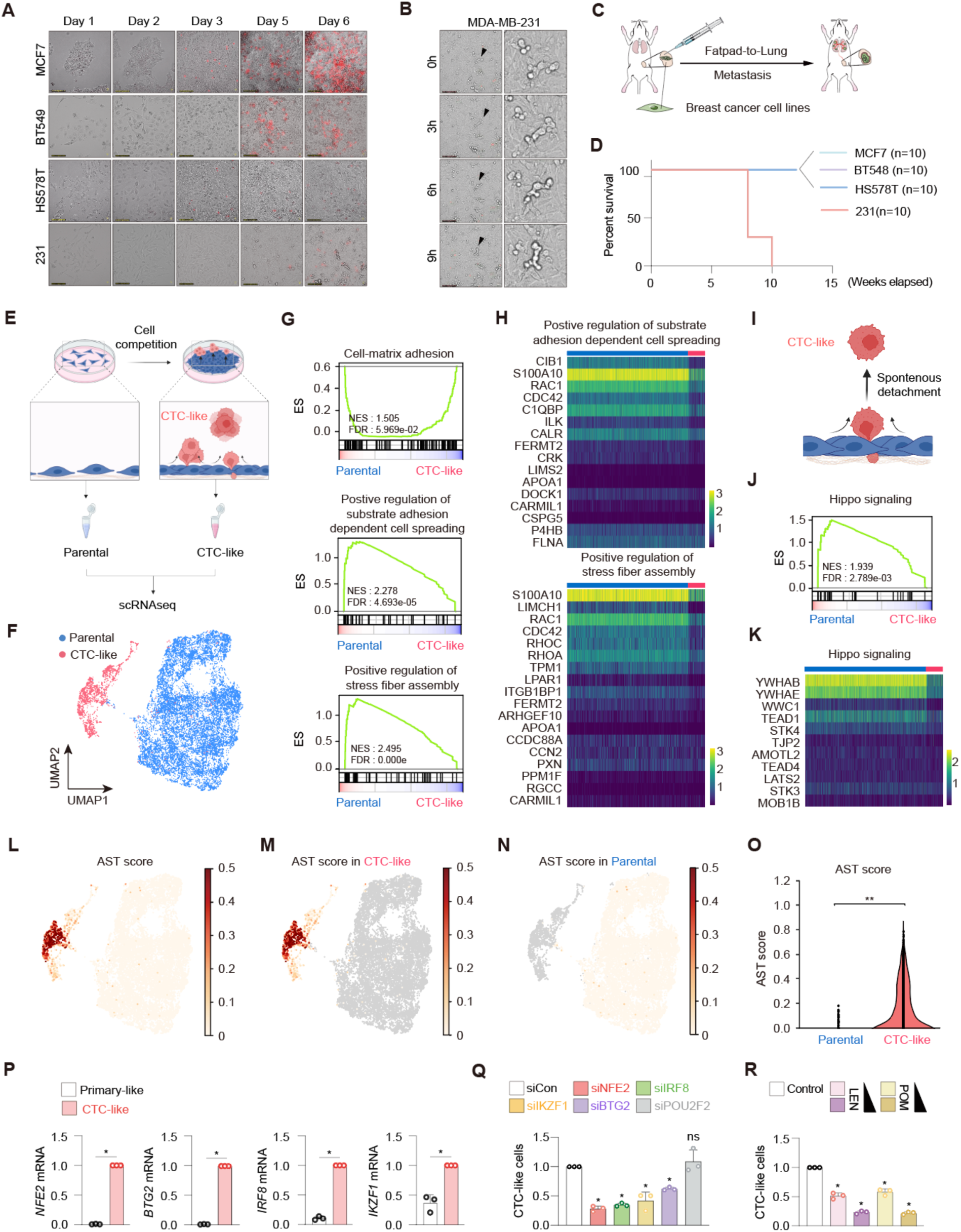
Cell competiton orchestrates spontaneous detachment of CTC-like cells driven by AST factors expression *in vitro*. **A**, Representative fluorescence images MCF7, BT549, Hs578T and MDA-MB-231 cells stained for viability with propidium iodide. Red, dead. Scale bars, 100 μm. **B**, Time-lapse microscopy images showing spontaneous cell detachment of MDA-MB-231 cells in the dissemination assay. **C**, Schematic overview of mammary fat pad xenografts of 4 different breast cancer cell lines. **D**, Kaplan-Meier survival plot showing the survival rates of mice injected with MCF7, BT549, Hs578T, or MDA-MB-231 cells. **E**, Schematic overview of the isolation and scRNA-seq analysis of parental cells and circulating tumor cells-like (CTC-like) cells from cell competition. **F**, Uniform manifold approximation and projection (UMAP) of scRNA-seq cells (13,918 cells) colored by parnetal (blue) and CTC-like cells (red). **G**, Gene set enrichment analysis (GSEA) results for cell-matrix adhesion, positive regulation of substrate adhesion dependent cell spreading, and positive regulation of stress fiber assembly gene sets within CTC-like cells. **H**, Heatmap showing differentially expressed genes (DEGs). DEGs were identified based on gene set enriched in GSEA result in (G). **I**, Schematic overview of spontaneously detached CTC-like cells from parental cells. **J**, Gene set enrichment analysis (GSEA) results for hippo signaling gene sets within CTC-like cells. **K**, Heatmap showing differentially expressed genes (DEGs). DEGs were identified based on gene set enriched in GSEA result in (K). **L**, UMAP of scRNA-seq cells showing AST score, calculated with Scnapy’s score_genes function, which reflects the normalized difference in mean expression between AST genes (IKZF1, NFE2, BTG2, and IRF8) and control genes. **M**, UMAP of scRNA-seq cells showing AST score in CTC-like cells. **N**, UMAP of scRNA-seq cells showing AST score in parental cells. **P,** Representative images of CTC-like MDA-MB-231 cell morphology and quantitative real-time PCR analysis of AST factor transcript levels in CTC-like cells. **Q**, The number of CTC-like cells for MDA-MB-231 cells following siRNA-mediated AST factor depletion. N = 3. **R**, The number of CTC-like cells for MDA-MB-231 cells following treatment with lenalidomide or pomalidomide. N = 3.

Next, to identify the molecular characteristics of cell competition-induced CTC-like cells, adherent parental cells and suspended CTC-like cells were subjected to single-cell RNA sequencing analysis (**Fig. 1E**). Interestingly, parental cells and CTC-like cells formed distinct clusters and GSEA analysis revealed that parental cells exhibited significant enrichment in gene sets associated with cell-matrix adhesion, substrate adhesion-dependent cell spreading, and stress fiber assembly compared to CTC-like cells, which indicates active downregulation of anchorage dependence is a hallmark of CTC-like cells (**Fig. 1F-G**). Notably, CTC-like cells exhibited a significant reduction in the expression of genes involved in focal adhesion formation, a process essential for CTCs to detach from the extracellular matrix and disseminate from the primary tumor (**Fig. 1H**). Recent reports have demonstrated the role of YAP1 and TEAD to resist mechanical pressures and confer growth advantage to tumors while the Hippo pathway, a negative regulator of YAP-TEAD, promotes cell-matrix dissociation and detachment of cancer cells from the primary tumors. Interestingly, our results show gene sets associated with Hippo signaling and the expression of core mediators of the Hippo pathway significantly reduced in CTC-like cells compared to parental cells (**Fig. 1I-J**). These results suggest that cell competition renders CTC-like cells competent to undergo spontaneous detachment from the matrix (**Fig. 1K**).

We firstly reported that solid tumor cells could spontaneously disseminate CTCs through reprogramming anchorage dependency via the aberrant expression of hematopoietic transcription factors referred to as adherent-to-suspension transition (AST) factors, IKZF1, IRF8, NFE2, and BTG2, which evoke global suppression of integrin-ECM expression, inhibition of YAP-TEAD activity, and induction of hemoglobin that resists anoikis. Thus, we sought to investigate whether cell competition reprograms loser cells with AST-related mechanisms. Surprisingly, cell competition-induced CTC-like cells recapitulated the aberrant induction of endogenous AST factors in MDA-MB-231, which recapitulate are previous findings in CTCs derived from mouse models and *de novo* metastatic breast cancer patients (**Fig. 1L**). scRNA-seq and qPCR results both indicate the significant expression of AST factors in CTC-like cells, but not in their adhesive counterparts (**Fig. 1M-P**). It is important to note that, in contrast to cell competition-induced CTC-like cells, trypsinization-induced passive detachment of cells fails to evoke the expression of AST factors (**Fig. S1B**). To verify whether the induction of AST factors is a prerequisite for the spontaneous detachment of CTC-like cells, we depleted each factor by siRNA and measured the viability of detached cells produced during cell competition. While combinatorial induction of four factors enhances the reprogramming efficiency, we found depletion of any single AST factor was sufficient to markedly suppress the formation of CTC-like cells (**Fig. 1Q**). Moreover, we found that pretreatment with thalidomide-derivatives such as lenalidomide or pomalidomide, which we previously identified as potential anti-metastatic agents that act by reducing the expression of IKZF1 in mouse breast cancer models ^7^, substantially suppressed the formation of CTC-like cells (**Fig. 1R**). These results demonstrate that cell competition reprograms anchorage dependency of adherent tumor cells to transit into CTC-like cells via upregulation of AST factors and related mechanisms.

### Cell competition-derived in vitro CTC-like cells reflect in vivo CTC characteristics

Next, we asked whether cell competition-derived *in vitro* CTC-like cells recapitulate *in vivo* CTCs disseminated from primary tumors in mouse models. To test this, we integrated scRNA-seq data obtained from parental and cell competition-derived CTC-like cells *in vitro* using the LM2 cell line, a derivative of MDA-MB-231 cells that efficiently forms lung metastasis, and compared it with scRNA-seq results from mouse tissue and liquid biopsy specimens that consist of orthotopically injected LM2-derived primary tumors and CTCs collected from a fat pad-to-lung metastasis model (**Fig. 2A**).

**Figure 2.**
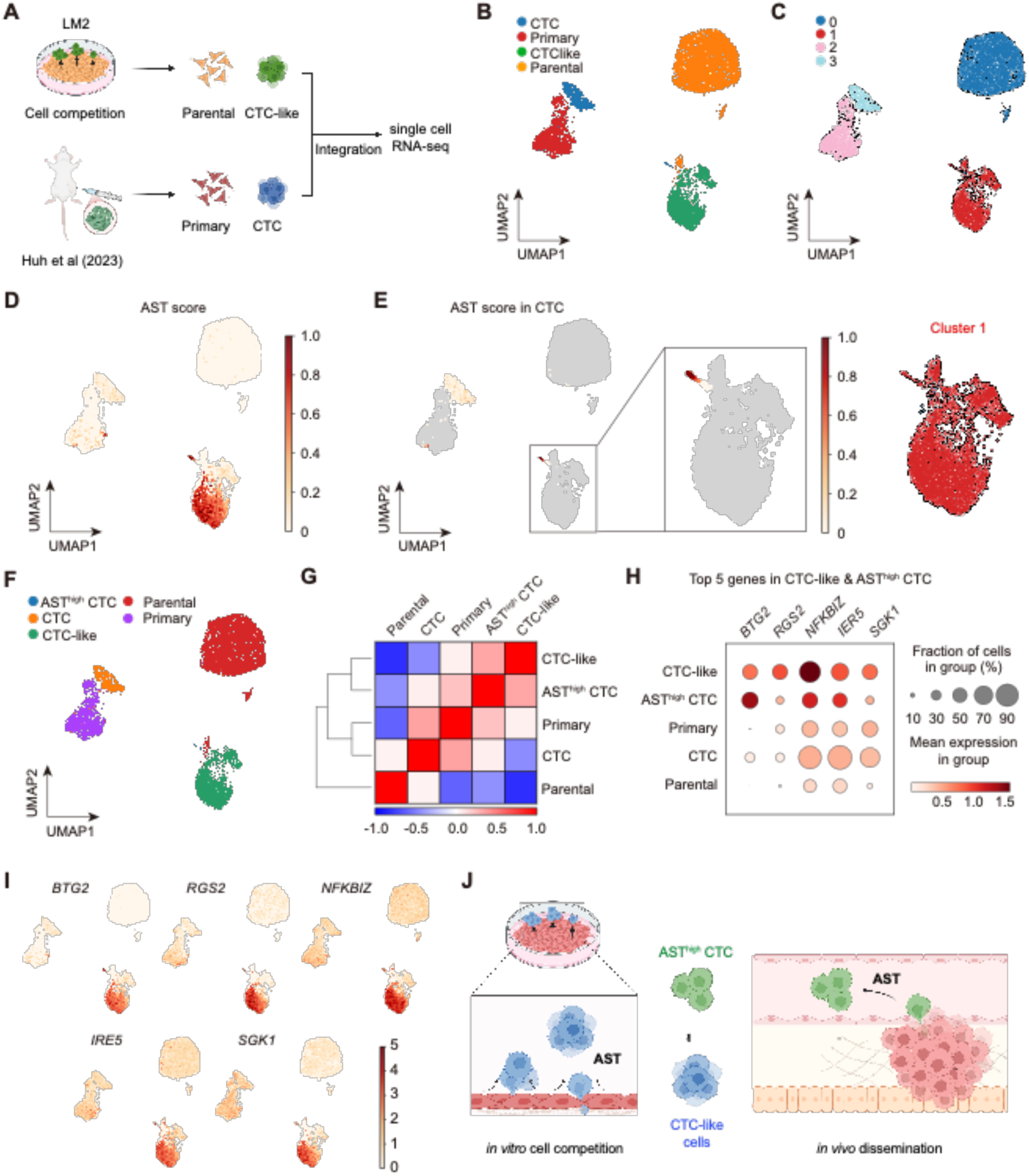
CTC-like cells reflect CTCs characterized by AST factor expression. **A,** Schematic overview illustrating the integration of scRNA-seq analysis of in vitro cell competition data (parental cells and CTC-like cells) with in vivo orthotopic fatpad-to-lung metastasis data (primary tumor and circulating tumor cells, CTCs). **B**, Uniform manifold approximation and projection (UMAP) of scRNA-seq cells (26,571 cells) colored by samples (CTCs, primary tumor, CTC-like cells and parental cells) **C**, Uniform manifold approximation and projection (UMAP) of scRNA-seq cells (26,571 cells) colored by leiden algorithm. **D**, Uniform manifold approximation and projection (UMAP) of scRNA-seq cells (26,571 cells) colored by normalized AST score, calculated with Scnapy’s score_genes function, which reflects the normalized difference in mean expression between AST genes (IKZF1, NFE2, BTG2, and IRF8) and control genes. **E**, Uniform manifold approximation and projection (UMAP) of scRNA-seq cells (26,571 cells) colored by normalized AST score in CTCs. **F**, Uniform manifold approximation and projection (UMAP) of scRNA-seq cells (26,571 cells) colored by annotated cell type (AST^high^ CTCs, CTC-like, CTCs, primary tumor, and parental cells). **G**, Pearson correlation analysis showing the transcriptional correation between annotated cell types in **F**. **H**, Dot plots showing the expression level of five top genes that is highly expressed in AST^high^ CTCs and CTC-like in annotated cell types in **F**. **I**, Uniform manifold approximation and projection (UMAP) of scRNA-seq cells (26,571 cells) colored by normalized expression of 5 top genes in **H**. **J**, Schematic model illustrating that in vitro CTC-like cells generated thorugh cell competition recapitulate in vivo AST^high^ CTCs.

In UMAP analysis, LM2 cells harvested from *in vitro* cell competition assay and *in vivo* mouse tissues were clustered into 4 groups (**Fig. 2B**). Surprisingly, we found that a portion of *in vivo* CTCs were clustered adjacent to the *in vitro* CTC-like cells and shared similar transcriptomic profiles based on Leiden algorithm (**Fig. 2C**). While most *in vivo* CTCs were located in Leiden cluster 3, a subpopulation of CTCs with significantly high AST score, which measures the expression level of AST factors, were grouped together with the CTC-like cell population in Leiden cluster 1 (**Fig. 2D-E**). This result suggests that the expression of AST factors may drive the transcriptomic similarities between *in vivo* CTCs and cell competition-derived CTC-like cells. Thus, among the in vivo CTCs harvested from mouse metastasis model, we annotated these cells as AST^high^ CTCs and investigated their characteristics compared to other groups (**Fig. 2F)**. Surprisingly, Pearson correlation analysis revealed that the transcriptome of *in vivo* AST^high^ CTCs exhibit the highest positive correlation with *in vitro* CTC-like cells, which was even stronger than their primary tumor counterparts (**Fig. 2G)**. GSEA analyses indicate AST^high^ CTCs exhibit impaired substrate adhesion, cell spreading, and Hippo signaling, which are hallmarks of CTCs in mouse models and patients (**Fig. 2H)**. Notably, AST^high^ CTCs and CTC-like cells shared common gene signatures related to downregulation of cell cycle, which resembles a feature of metastasis-initiating cells that enter slow-cycling states for dormancy to survive the stresses of metasatic processes (**Fig. 2I-J)**. We further identified the top five genes strongly expressed in both CTC-like cells and AST^high^ CTCs, including BTG2, RGS2, NFKBIZ, IRE5, and SGK1, which have shared functional roles in stress-adaptive quiescence that promotes CTC survival in hostile niches (**Fig. 2H-J)**. This result strongly demonstrates that CTC-like cells generated through cell competition physiologically recapitulate disseminated CTCs from the primary tumor that have undergone AST-mediated reprogramming (**Fig. 2M)**.

### Oscillatory dynamics of AST factors confer anchorage plasticity of CTCs

Characterization of cell competition-induced CTCs provides the basis to further investigate the significance of AST factor plasticity in mouse models and human patients. To examine the anchorage plasticity of CTCs, the loser cells or CTC-like cells were subsequently replated after dissemination assay. Surprisingly, we found that CTC-like cells lost their anchorage independence and readily colonized the culture dish to resume proliferation as adherent cells. We refer to these newly adherent cells as mets-like cells because they simulate the latent phase of metastasis where CTCs regain adhesive properties to reattach to the distant organs (**Fig. 3A**). Surprisingly, the spontaneously detached CTC-like cells showed a significant reduction in AST factor expression, which promoted their spontaneous detachment as mets-like cells, having undergone the suspension-to-adherent transition (SAT) (**Fig. 3B**). These results show that AST factor dynamics in CTC-like cells undergoing from AST to SAT transition correlate with the anchorage plasticity of solid tumor cells in a way that is quite distinct from the primary-like or met-like cells. Our findings demonstrate AST factors as anchorage-dependent, mechanosensitive transcriptional regulators, highlighting the need for further investigation into the molecular mechanisms by which mechanical cues modulate their transcriptional activity in solid tumor cells.

**Figure 3.**
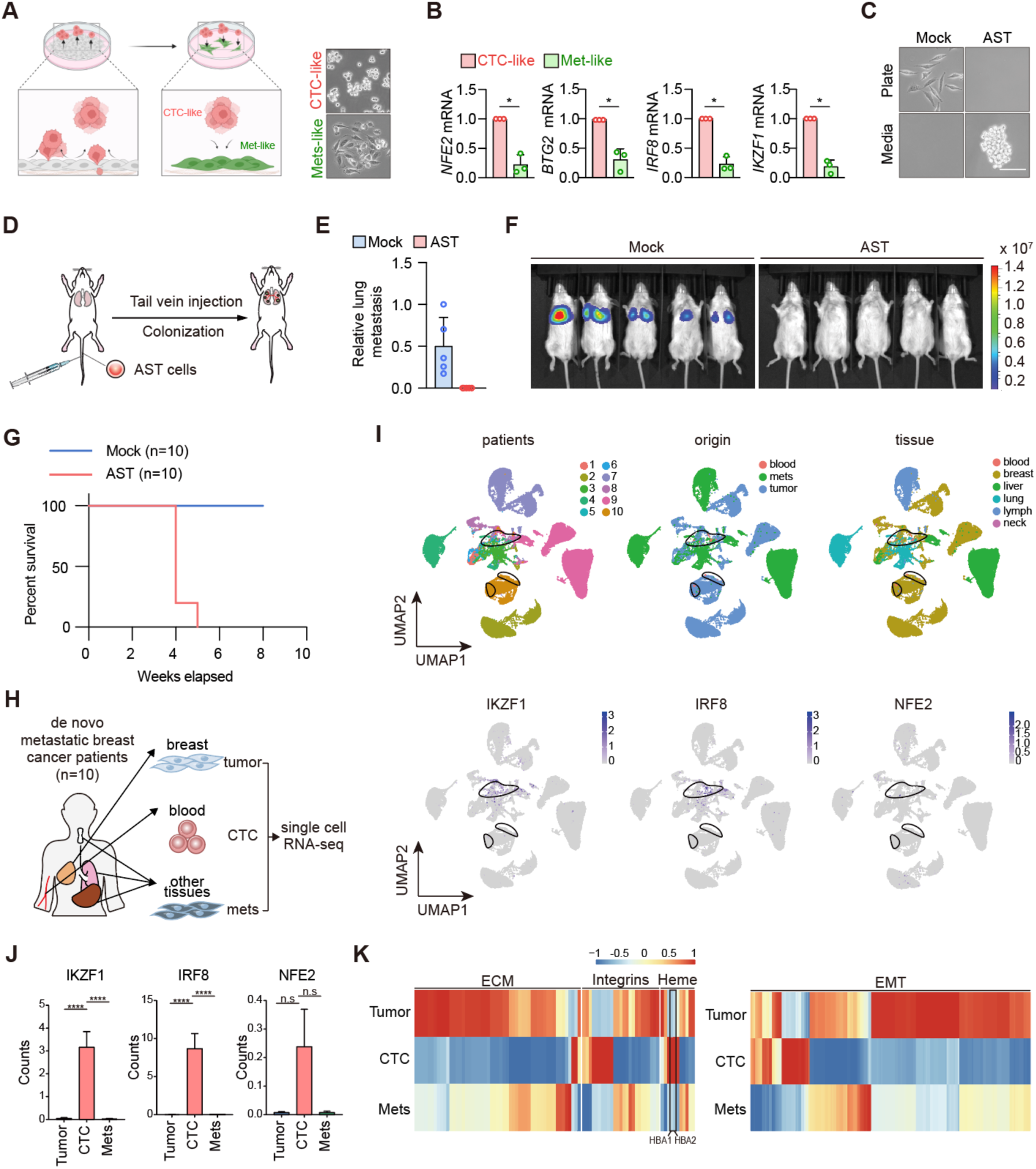
CTC-like cells and CTCs confer anchorage dependency plasticity accompanied by AST factors oscillation. **A,** Schematic overview of CTC-like cells and mets-like cells, and representative images of re-attached MDA-MB-231 cell (met-like cells) morphology. **B**, Quantitative real-time PCR analysis of AST factor transcript levels in met-like cells compared to CTC-like cells. **C**, Representative images of MDA-MB-231 derivates LM2-mock or AST cell (LM2 cells expressing AST factors) morphology on a culture plate or in media. **D**, Schematic overview of tail vein injections with LM2-mock or AST cells. **E**, Bioluminescence images of lung metastases after LM2 tail vein injections. **F**, Lung metastases quantified by lung bioluminescence intensity for mock versus AST cells. **G**, Kaplan-Meier survival plot showing the survival rates of mice injected with mock or 3AST cells. N = 10. **H,** Schematic overview of the isolation and scRNA-seq analysis of primary tumor cells, circulating tumor cells (CTCs), and metastatic tumor cells from de novo metastatic breast cancer patients (n = 10). CTCs were isolated using HDM chips. **I,** Uniform manifold approximation and projection (UMAP) of scRNA-seq cells (n = 10; 54,866 cells) colored by patient, origin, tissue, and normalized gene expression of the AST factors IKZF1, IRF8, and NFE2. Black border lines, area in which the CTCs are located. **J,** Dynamite plots showing the expression of (left) IKZF1, (middle) IRF8, and (right) NFE2 in primary tumor cells, CTCs, and metastatic tumor cells. ****, P < 0.0001; n.s, not significant (P > 0.05). **K**, Heatmaps showing normalized expression of four gene sets—ECM, integrins, antioxidant defense genes, and EMT-related genes—in primary tumors, CTCs, and metastases. The average normalized expression values for cells within each group are displayed as scaled values. Black-bordered box, HBA1 and HBA2.

Building on our prior findings that AST factor induction facilitates CTC dissemination, we next investigated whether repression of AST factors is a prerequisite for the reattachment and metastatic colonization of CTCs *in vivo*. To this end, we established CTC-like LM2 cells by stably expressing the AST factors (**Fig. 3C**).^8^ Unlike mock-transduced adherent control cells, LM2-AST cells elicited spontaneous detachment and resistance to anoikis. To assess whether sustained AST factor expression with impaired plasticity could hinder metastatic colonization, we performed tail vein injections of LM2-control and LM2-AST cells into NOD/SCID gamma (NSG) mice and monitored lung colonization (**Fig. 3D**). Remarkably, while LM2-control cells efficiently formed lung metastases, constitutive expression of AST factors in LM2-AST cells significantly impaired lung colonization and extended overall survival **(Fig. 3E–G)**. These findings suggest that while AST factor induction promotes CTC dissemination, their subsequent repression is essential for the successful metastatic colonization of CTCs. Collectively, our data indicate that the dynamic oscillation of AST factors, transient induction termed AST during the primary-to-CTC transition followed by repression termed SAT during the CTC-to-metastasis transition, is a critical determinant of the metastatic cascade.

Next, we asked whether Adherent-to-Suspension Plasticity (ASP) represents a clinically relevant mechanism that dynamically regulates the anchorage dependency of CTCs during the metastatic cascade in patients. To assess the clinical relevance of ASP in human patients, we prospectively recruited 10 newly diagnosed *de novo* metastatic breast cancer patients and collected matched specimens of primary breast tumors, peripheral blood, and metastatic lesions from each patient **(Table S1, Fig. 3H)**. This cohort represents one of the most comprehensive biobanks in breast cancer integrating both tissue and liquid biopsy specimens from the same individuals, offering invaluable opportunities to investigate metastatic progression at the single-cell level^1–4^. CTCs were enriched from whole blood using high-density microporous (HDM) chips via size-based exclusion, and tissue-derived cells and CTCs were processed for single-cell RNA sequencing (scRNA-seq) analysis^9^. We identified multiple major cell types, including epithelial tumor cells, fibroblasts, mesenchymal stem cells, and various immune populations such as myeloid and lymphoid subsets (**Fig.S2A-B**). Tumor cells were identified by co-expression of EPCAM and breast cancer-specific cytokeratins (KRT5, KRT8, KRT9, KRT17, or KRT18). Importantly, UMAP projection not only revealed patient- and tissue-specific clustering, but also highlighted a central, mixed cluster comprising cells from multiple patients and compartments, in which the majority of CTCs were located (**Fig.3I**). Strikingly, this central cluster exhibited elevated expression of AST factors, including IKZF1, NFE2, and IRF8. Quantitative analysis revealed that AST factor expression was significantly higher in CTCs compared to both primary and metastatic tumor cells, with IKZF1 and IRF8 exceeding statistical thresholds **(Fig. 3J, Fig. S2C)**. Notably, this increase was consistent across different breast cancer subtypes and metastatic organ sites, supporting the clinical relevance of ASP as a conserved mechanism in CTC biology. These findings establish the upregulation of AST factors as a hallmark of anchorage plasticity during metastatic dissemination in human breast cancer.

Extending upon our previous gene set enrichment analysis (GSEA) comparing primary tumors and CTCs, we observed a significant enrichment of focal adhesion and extracellular matrix (ECM) organization pathways in tissue-derived tumor cells relative to CTCs (**Fig.3K**)^7^. Specifically, focal adhesion-related gene expression was markedly elevated in both primary and metastatic tumor samples compared to CTCs, while ECM organization pathways were predominantly enriched in metastatic lesions (**Fig. S2D**). In contrast, CTCs exhibited downregulation of genes encoding ECM components, integrins, and EMT-related genes consistent with their anchorage-independent phenotype. Concurrently, CTCs showed a significant upregulation of hemoglobin genes HBA1 and HBA2, which have been implicated in neutralizing reactive oxygen species (ROS) and promoting resistance to anoikis (**Fig. 3K**)^7^. These transcriptomic changes in matched tissue and liquid biopsy specimens from metastatic cancer patients support the model that AST factor dynamics regulate anchorage plasticity through transcriptional reprogramming of integrin/ECM-associated pathways and oxidative stress defense in patients. Collectively, out results demonstrate the clinical relevance and mechanisms of Adherent-to-Suspension Plasticity (ASP) in mediating the anchorage dependance of CTCs in metastasis.

### Adherent-suspension plasticity in metastatic breast cancer patients

To characterize the transcriptional dynamics underlying CTC formation and their relationship with solid tumor cells, we further analyzed scRNA-seq database on matched primary tumors, CTCs, and metastatic lesions from patients with *de novo* metastatic breast cancer. We first conducted pseudotime and UMAP projection analyses to visualize cell state transitions across the metastatic cascade (**Fig. 4A**). While cells clustered based on tissue origin, cancer stage, and patient identity, it is important to note that a distinct central region emerged, where primary, metastatic, and CTC populations overlapped in pseudotime space (**Fig. 4B**). Based on gene expression similarity and reduced transcriptomic variance, we identified eight clusters (9, 11, 12, 13, 15, 17, 20, and 24) that define the central region, which comprised 8,823 solid tumor cells (3,695 primary and 5,128 metastatic) and 39 CTCs (**Fig. 4C-D, Fig. S3A-B**).

**Figure 4.**
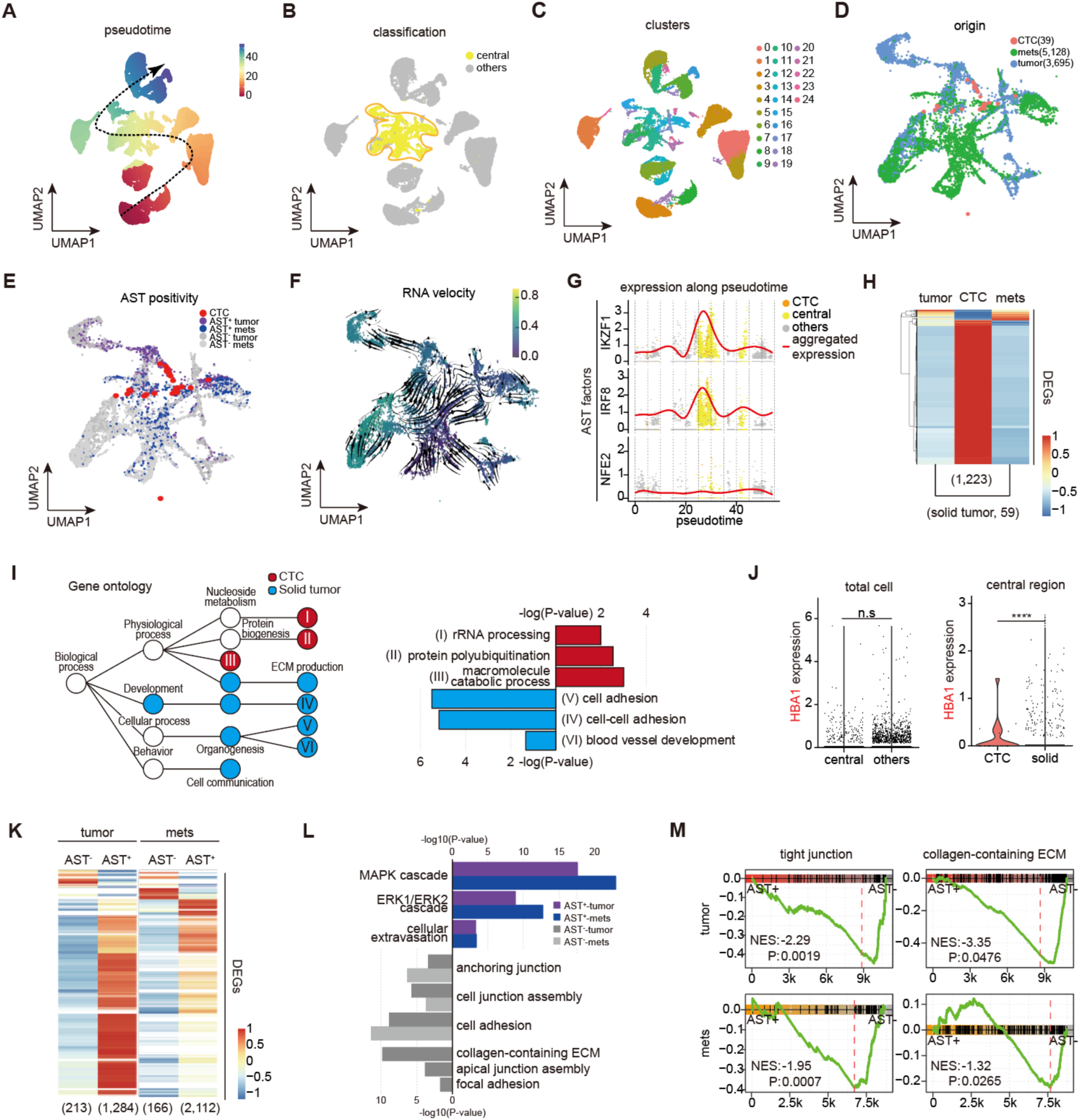
AST plasticity in matched single cell RNA-seq samples from *de novo* breast cancer patients. **A-C**, UMAP plot showing pseudotime (A), a division between the central region containing complex clusters with CTCs and others (B), and clusters calculated at a resolution of 0.5 (C). Dotted arrow, flow of pseudotime for the normalized gene expression of the AST factors IKZF1, IRF8, and NFE2. **D-F**, UMAP of the central region colored according to origin (D), AST positivity (E), and RNA velocity (F). **G**, Pseudotime expression plots showing AST factors. X-axis, pseudotime; y-axis, expression levels; gray dots, others; yellow dots, central region; orange dots, CTCs; red line, linear expression trend using generalized additive modeling (GAM). **H**, Heatmap showing differentially expressed genes (DEGs). DEGs were identified based on comparisons between solid tumor cells and CTCs within the central region and selected with a threshold p-value < 0.05. There were 1,223 CTC DEGs and 59 solid tumor DEGs. **I**, Gene ontology tree showing the CTC-specific gene set and the solid tumor-specific gene set with a bar plot showing the gene names and the p-values corresponding to the circles marked with roman numerals. Colored circles, statistically significant, FDR corrected P < 0.01. **J**, Violin plots with dots showing HBA1 expression. The first plot compares the central region with other groups, while the second plot compares CTCs with solid tumor cells within the central region. Yellow, central region; gray, others; orange, CTCs within the central region; turquoise, solid tumor cells within the central region; ****, P < 0.0001; n.s, not significant (P > 0.05). **K**, Heatmap showing DEGs. In a comparison of cells with and without AST factor expression in both primary and metastatic tumors, the DEG thresholds were defined as an FDR corrected q-value < 0.001 and an absolute fold change > 2. **L**, Bar plot showing ontology analysis results for the AST-positive and AST-negative gene sets identified in (**K**). Purple bars, AST-positive gene sets in primary tumors; blue bars, AST-positive gene sets in metastatic tumors; gray bars, AST-negative gene sets in primary tumors; light gray bars, AST-negative gene sets in metastatic tumors. **M**, Gene set enrichment analysis (GSEA) results for tight junction and collagen-containing ECM gene sets within both primary and metastatic tumors.

Next, we further analyzed the origin of these central region cells and observed that the majority of CTCs were localized in this region, which shared the pseudotime trajectories with primary and metastatic tumor cells (**Fig. 4D**). We then examined AST factor expression in cells located within the central region. Surprisingly, AST-positive cells (AST^+^), including CTCs and a subset of solid tumor cells were enriched in this region (**Fig. 4E**). RNA velocity analysis further supported this transitionary state, indicating dynamic transcriptional activity toward a suspension phenotype (**Fig. 4F**). Pseudotime expression plots revealed the plasticity of AST factors such as IKZF1 and IRF8, which were significantly upregulated in intermediate states that correspond to the central region, compared to early and later stages. These findings are consistent with the oscillatory dynamics of AST factors shown in dissemination assay-derived CTC-like cells (**Fig. 4G**). Moreover, these results suggest that the transcriptional profile of solid tumor cells from primary and metastatic lesions within the central region resembles gene signatures of CTCs, indicating an intermediate state, not yet fully transformed into CTCs.

To further dissect the molecular basis of anchorage plasticity during the metastatic cascade, we compared CTCs with matched solid tumor cells within the central region and identified 1,223 CTC-specific and 59 solid tumor-specific genes (**Fig. 4H, S3C**). Gene ontology analysis revealed that CTC-specific genes were strongly enriched in rRNA processing and ribosome biogenesis pathways, consistent with the elevated protein synthesis required for survival and adaptation of disseminated CTCs (**Fig. 4I**). Notably, hemoglobin genes such as HBA1, which provide antioxidant defense and promote anoikis resistance, were markedly upregulated in CTCs (**Fig. 4J**). In contrast, solid tumor–specific genes were enriched for pathways related to cell–matrix adhesion and extracellular matrix (ECM) organization (**Fig. 4I**), reflecting their anchorage-dependent state. Together, these results indicate that CTCs transcriptionally reprogram toward an anchorage-independent phenotype, supported by ribosome biogenesis and oxidative stress resistance, whereas solid tumor cells retain adhesion and ECM-related programs underscoring the role of AST factor-driven anchorage plasticity in mediating the CTC transition.

Pseudotime analysis further delineated the emergence of CTC-like states within solid tumor specimens by showing that while the majority of tumor cells lacked AST factor expression (AST-negative), a subset located at pseudotime positions proximal to CTCs displayed robust AST factor expression (AST-positive) (**Fig. 4E–F**). We conducted DEG analyses across both primary and metastatic tumors to investigate the differences between AST-positive and AST-negative solid tumor cells. This allowed us to identify 2,112 AST-positive-specific genes and 166 AST-negative-specific genes in the primary tumors versus 1,284 AST-positive-specific genes and 213 AST-negative-specific genes in the metastatic tumors (**Fig**. **4K**). Underscoring the similarity of the two conditions, we found a considerable overlap of 997 AST-positive-specific genes and 45 AST-negative-specific genes that were shared between the primary and metastatic tumors. A GO analysis of these AST-positive-specific genes revealed enrichment in the MAPK-ERK1/2 cascade, which is associated with antioxidant expression, anoikis resistance, and metalloproteinase-mediated ECM degradation (**Fig**. **4L**) ^10–13^. In the AST-positive cells, we also identified genes related to extravasation, which suggests that, like CTCs, these cells possess increased invasive properties. In contrast, GO analysis of the AST-negative-specific genes revealed enrichment in pathways related to anchoring junctions, ECM, and focal adhesions, which promote cell adhesion and structural integrity (**Fig**. **4L**). GSEA analysis confirmed that AST-negative cells in both primary and metastatic tumors showed significant enrichment in gene sets related to tight junctions and ECM organization (**Fig**. **4M**). This further supports the notion that AST-negative cells are characterized by strong adhesion properties. In contrast, AST-positive cells, particularly in the central region, co-expressed genes involved in the MAPK cascade, ERK1/2 cascade, and extravasation. Thus, some solid tumor cells exhibit an expression pattern like that of CTCs, indicating the presence of cells with CTC-like characteristics within the solid tumor population.

Finally, we analyzed RNA velocity to clarify the correlation of AST factor expression dynamics with the expression of integrin/ECM-related, antioxidant-related, and EMT-related genes (**Fig. 4F, S3D**). Although we found no correlation between AST factors and EMT-related genes, there was a correlation between EMT-related genes and ECM-related genes, which correlates with our previous study that AST is independent of EMT.

## Discussion

Cell competition in tumors is a system that leverages the heterogeneity of cancer cells in stressful microenvironments, such as high cell density, to select for cells that contribute to tumor survival and progression. During this process, less fit cells either undergo apoptosis or detach from the existing colonies. Recent studies have indicated that cells with metastatic traits can lose their competitive advantage within primary tumors and detach. In our study, we found that cells detached from the primary tumor due to cell competition acquire anchorage independence via AST factors, and are transcriptionally similar to circulating tumor cells (CTCs) expressing AST factors. This suggests the potential for conducting in vitro studies on CTCs using cell competition assays with metastatic cells. Previous attempts have aimed to study CTCs at the molecular level in vitro, including efforts to establish ex vivo CTC cell lines from patient samples. Furthermore, suspension assays, which assess anchorage-independent growth, have been widely used to investigate the anoikis resistance of CTCs. However, because this assay use trypsinized cells, they cannot mimic the spontaneous process by which CTCs disseminate *in vivo*, nor can they generate the exceedingly rare population of CTC-like cells that can freely detach from adherent culture conditions. Therefore, the CTC-like cells identified through cell competition will shed light on the overall understanding of the mechanical modulation involved in the dissemination and survival of AST-associated CTCs, paving the way for the discovery of potential anti-metastatic agents.

Our study highlights the need to distinguish anti-metastatic strategies from traditional cancer therapies that promote competitive cells towards a less fit direction within the primary tumor’s cell competition context. For instance, cancer therapies targeting Hippo pathway effector proteins such as TEAD or YAP suppress tumor proliferation by inhibiting the competitive advantage of YAP^hi^ or TEAD^hi^ cells. However, based on our findings that less fit CTC-like cells or CTCs dissociated from the parental population are TEAD^low^, targeting TEAD could inadvertently increase the less fit cell population, potentially promoting metastasis. Therefore, anti-cancer therapies and anti-metastatic agents should be developed with a more nuanced understanding of how shifts in the dominance between competitive and less fit cells in the context of cell competition within primary tumors can influence tumorigenesis and metastasis, respectively. This approach would ensure that anti-cancer therapies do not inadvertently promote metastasis, and anti-metastatic agents do not compromise tumor progression.

We demonstrated that, while AST factors facilitate the initial detachment and anoikis resistance of CTC-like cells, their downregulation is essential for the reattachment and successful colonization of cancer cells at distant metastatic sites. This indicates that metastatic success depends on the ability of cancer cells to adapt their anchorage dependency to each stage of metastasis. Suspension characteristics mobilize CTCs into the bloodstream, whereas reattachment and colony formation necessitate a reversion to adherent traits within the microenvironment of secondary sites. These insights challenge the conventional understanding of metastasis as a linear progression and instead propose a model in which anchorage plasticity plays a central role in cancer cells to spread during metastasis. ASP thus represents a novel cellular plasticity mechanism analogous to epithelial-mesenchymal plasticity (EMP), another well-established process that facilitates metastasis, however, limited to the transition between two different adhesive cell types. How cancer cells orchestrate ASP and EMP during each stage of the metastatic cascade warrants further investigation.

Our findings underscore the pivotal role of ASP in the metastatic cascade, as demonstrated by scRNA-seq analysis of paired samples from *de novo* metastatic breast cancer patients, encompassing matched primary tumors, CTCs, and metastatic lesions. This dataset offers invaluable insights into tumor progression, adaptability, and heterogeneity across metastatic stages in individual patients. The aggregation of CTCs from different patients within the central region of the UMAP signifies that CTCs exhibit similar transcriptomic profiles despite their origin. Importantly, the significant expression of AST factors not only in CTCs but also in their intermediate solid tumor counterparts from pseudotime analysis indicate that anchorage plasticity from AST to SAT is an adaptive and continuous process throughout the metastatic cascade. The consistency of these trends across tumor subtypes and metastatic locations underscores the broad significance of AST factors as theragnostic biomarkers of metastasis. To elucidate the role of tumor microenvironment as an upstream mediator of AST factors, the spatiotemporal expression patterns of AST factors using matched longitudinal specimens warrants further investigation.

Our study provides the rationale for clinical implementation of therapeutic strategies that target AST factors and related mechanisms. For example, therapies designed to inhibit the transition from adherent-to-suspension states, referred to as anti-disseminating agents, could potentially be used as neoadjuvant therapies or alone to impede the initial detachment of cancer cells from the primary tumor, thereby reducing the number of CTCs and the likelihood of metastatic spread. Conversely, strategies that prevent the reattachment of CTCs at secondary sites could hinder metastatic colonization, offering a two-pronged approach to tackle metastasis. A deeper exploration into the genetic and epigenetic regulation of AST factors could yield additional mechanistic insights into ASP. Understanding how AST factors are modulated during different stages of metastasis, and the interplay between ASP and other forms of cellular plasticity, such as EMP, warrants further investigation to delineate how these processes collectively contribute to the completion of metastasis.

Together, our findings highlight the unprecedented role of ASP in breast cancer metastasis. ASP provides a novel theoretical basis in cancer metastasis and suggests AST factors as promising therapeutic targets to tackle the metastatic spread of solid tumor cells. Clinically, moving beyond the primary tumor-centric point-of-view, our data expand the prevailing cancer treatment paradigm toward direct intervention within the metastatic processes. The availability of cell-based platforms that measure ASP efficiency and multi-omics database of matched longitudinal patient samples could facilitate future research opportunities, including individualized metastasis prediction models, hence augmenting the potential for developing effective anti-metastatic therapies ^14^.

## METHODS

### DNA constructs

AST genes were tagged with V5 and subcloned into the pENTR4 vector (Addgene, 17425 and 17423) using standard molecular cloning techniques. The resulting subcloned vectors were then recombined with the destination vector pLentiCMV (Addgene, 17293) using LR recombinase (Invitrogen, 1179019), generating the final lentiviral expression vectors.

### Cell culture

Cells were maintained in a 37 °C humidified incubator with 5% CO_2_. HEK293T, MCF7, HS578T and MDA-MB-231 cells were cultured in DMEM (Hyclone, SH30022.01). BT549 cells were cultured in RPMI (Hyclone, SH30027.01). Culture media were supplemented with 10% FBS (Hyclone, SV30207.02) and 1% penicillin/streptomycin (Invitrogen, 15140122).

### Cell competition assay

Cell competition assays were performed using MDA-MB-231, MCF7, BT549, and HS578T cells, with each cell line cultured in its own respective growth media. For the assay, the cells were seeded on normal plates at 80% confluency on the first day and then maintained for 3 days without any media changes. After 3 days, the cells surpass 100% confluency, resulting in very high cell density. After the old media were replaced with fresh media, the cells were cultured for an additional 3 days at 100% confluency. We noticed MDA-MB-231 cells began to round up and detach from the plate beginning on the second day after the media change. By the third day, we observed a higher number of CTC-like cells. These detached CTC-like cells were harvested by collecting the media, followed by centrifugation to isolate the cells. For MCF7 cells, despite the media change, the cells began to die after 3 days. Although BT549 and HS578T cells did not exhibit cell death, they did cease proliferating.

### Viral infection

To generate lentiviral particles, HEK293T cells were transfected with plasmids encoding pMD2G and psPAX2, along with the lentiviral vectors containing the desired constructs. Transfections were performed using Polyplus Reagent (Merck) according to the manufacturer’s protocol. At 48 hours post-transfection, media containing the viral particles were harvested from the transfected cells. The harvested media were then filtered through a 0.45 µm filter to remove cellular debris. To enhance infection efficiency, the filtered media were supplemented with 8 mg/ml polybrene. The transduced cells were infected with the lentiviral particles by incubating them with the filtered media. After 24 hours of infection, the media were changed, and the cells were incubated for another 24 hours. Then, transduced cells were selected using puromycin. The selection process allowed for the enrichment of cells that successfully integrated the lentiviral vectors and expressed the desired genes or markers.

### Cell viability assay

*In vitro* cell viability was assessed using the CellTiter Glo® 2.0 cell viability assay (Promega, G9241) and the LIVE/DEAD® cell imaging kit (ThermoFisher, R37601) according to their manufacturers’ protocols. For the suspension assay, cells were plated on Corning® Costar® Ultra-Low Attachment Multiple-Well Plates (Merck, CLS3471).

### Quantitative real-time PCR analysis

Cellular RNA samples were collected via RNA extraction using the RNeasy Plus mini kit (QIAGEN, 74136). These RNA samples were then reverse transcribed into complementary DNA (cDNA) using iScript reverse transcriptase (Bio-Rad, 1708891). The KAPA SYBR FAST qPCR kit (Kapa Biosystems, KK4605) and the 7300 real-time PCR system (Applied Biosystems) were used for quantitative real-time PCR (qRT-PCR). The following primer sequences were used for qPCR: IKZF1, 5’-TTTCAGGGAAGGAAAGCCCC-3’ (forward) and 5’- CTCCGCACATTCTTCCCCAT-3’ (reverse); NFE2, 5’-ACAGCTGTCCACTTCAGAGC-3’ (forward) and 5’-TGAGCAGGGGCAGTAAGTTG-3’ (reverse); IRF8, 5’- AGCATGTTCCGGATCCCTTG-3’ (forward) and 5’-CGGTCCGTCACTTCCTCAAA-3’ (reverse); and ACTB, 5’-GCCGACAGGATGCAGAAGGAGATCA-3’(forward) and 5’- AAGCATTTGCGGTGGACGATGGA-3’(reverse).

### siRNA transfection

A transfection mix was prepared by combining 100 µl of Opti-MEM™ medium (ThermoFisher, #31985070), 12 µl of RNAi Max transfection reagent (Invitrogen, 13778150), and 2 µl of 20 µM siRNA. The following siRNAs were used for transfection: non-silencing (NS) siRNA (Dharmacon siGENOME non-targeting control pool #D-001206-13-200), siIKZF1 (Dharmacon siGENOME SMARTPool #M-019092-01-0010), siNFE2 (Dharmacon siGENOME SMARTPool #M-010049-00-0010), siIRF8 (Dharmacon siGENOME SMARTPool #M-011699-01-0010), and siPOU2F2 (Dharmacon siGENOME SMARTPool #M-019690-01-0010).

### Mouse model experiments

All animal experiment protocols in this study underwent a thorough review and received approval from the Yonsei University Institutional Animal Care and Use Committee (IACUC). All procedures involving animals adhered to the Guidelines for the Care and Use of Laboratory Animals. The NOD SCID Gamma (NSG) mice used for orthotopic injection experiments were obtained from JA Bio. For the orthotopic mouse model experiments, mammary fat pads of 7-week-old female NSG mice were orthotopically injected with 1x10^6^ MCF7, BT549, HS578T, or MDA-MB-231 cells. For the mouse tail vein injection experiments, the tail veins of 7-week-old female NSG mice were injected with 1x10^5^ MDA-MB-231 derivative LM2 cells. At week 4, the mice were euthanized for bioluminescence imaging using the In Vivo Imaging Spectrum (IVIS) device (PerkinElmer, CLS136331 IVIS Lumina LT Inst, Series III, 120 V). All *in vivo* experiments were carried out using mice that had received the same treatment. The NSG mice used in these experiments possessed an identical genotype and genetic background.

### Statistical analyses

All quantitative data were obtained from at least three independent biological replicates. Data are presented as means ± standard deviation (SD) unless otherwise noted in the figure legends. Statistical differences between two groups were examined using two-tailed, unpaired Student’s t-tests or one-way analysis of variance (ANOVA) with Bonferroni corrections for multiple comparisons. Statistical tests were performed using the GraphPad Prism 9.0 software (GraphPad Software, CA, USA). Two-sided p-values of less than 0.05 were considered significant. No statistical methods were used to predetermine sample size. Sample size was based on previous experience with experimental variability. Blinding was performed wherever possible during all sample analysis by coding sample identity during data collection and having the analysis performed by an observer without knowledge of or access to the experimental conditions.

### Bioinformatics of primary tumor cells, CTCs, and metastatic cancer cells

The illumina outputs for human samples were mapped to the human reference genome (GRCh38) using Cell Ranger 7.1.0 (10X Genomics). Gene-wise read counts for genes with at least 500 reads were exported from Cell Ranger to the Matrix Market format and imported into R with Seurat’s Read10X function. An average of > 135 million reads from > 13,000 cells were obtained for each sample. Each 10X library was individually checked for quality, and the cells were filtered according to the same criteria used in our previous study to ensure good gene coverage ^7^. After quality control filtering, a total of 310,189 cells remained, with 69,612 cells from the primary tumor site, 171,504 cells from the blood, and 79,073 cells from the metastatic site (**Table 1**). Samples were combined via an algorithm implemented in Seurat. The default Seurat function settings were used, and 1:20 principal component dimensions were used for all dimension reduction and integration steps. The cluster resolutions were set to 0.5, unless otherwise stated. The RunUMAP random seed was set to 1,000 to ensure reproducibility. Marker genes for cell clusters were identified using Seurat’s FindMarkers function with the default settings. Cell type annotation was performed using the SingleR package (v2.6.0) with the HumanProteinAtlas database from the celldex package (v1.14.0). Cells co-expressing EPCAM and cytokeratins such as KRT5, KRT8, KRT9, KRT17, or KRT18 were designated as cancer cells. A total of 54,867 cells were used for the analysis, including 26,227 cells from the primary tumor site, 50 cells from the blood, and 28,590 cells from metastatic sites (**Table 1**). The cells designated as cancer cells were subsequently extracted and reanalyzed beginning with the normalization process. DEGs were selected with thresholds for the FDR corrected p-value < 0.001 and an absolute log_2_ fold change ≥ 1. GO analysis was performed using the gseGO function from clusterProfiler (v3.0.4). GSEA plots were generated from the DEG data using the gseaplot function. Pseudotime analysis was performed using the monocle package (v3.0) with normalized data, UMAP coordinates, and cluster information obtained from the initial Seurat analysis. The calculation of gene set scores along pseudotime was conducted for each cell using the AddModuleScore function in Seurat. All plots were exported to the PDF format using R and their dimensions were adjusted in Illustrator.

## Declarations

### Ethical approval

Ethical approval for the use of human subjects was obtained from the Institutional Review Board of Gangnam Severance Hospital, Yonsei University, Seoul, Republic of Korea (3-2021-0236) in compliance with the ethical guidelines of the 1975 Declaration of Helsinki, and informed written consent was obtained from each patient. Collection of human breast cancer samples were obtained from consenting patients in accordance with guidelines and regulations approved by the Institutional Review Board of Gangnam Severance Hospital, Yonsei University Health System (3-2021-0236). The animal experiment protocols were reviewed and approved by the Yonsei University Institutional Animal Care and Use Committee (IACUC).

### Fundings and acknowledgments

This work was supported by grants from the National Research Foundation of Korea (RS-2024-00509461 to H.W.P), by the Brain Korea 21 FOUR Program (to H.D.H., Y.S., H.L., B.J.H., W.Y.P., J.O., and D.K.L.)

### Authors’ contributions

H.D.H., Y.S., and J.H.K. performed most of the experiments. H.L., B.J.H., W.Y.P. performed drug treatment and imaging. J.O., and D.K.L. performed transcriptome analyses. S.P. and J.C. performed CTC isolation and scRNAseq. J.J., H.Y.G., and H.W.P. designed the study and wrote the manuscript.

---

**Supplementary Figure 1.**
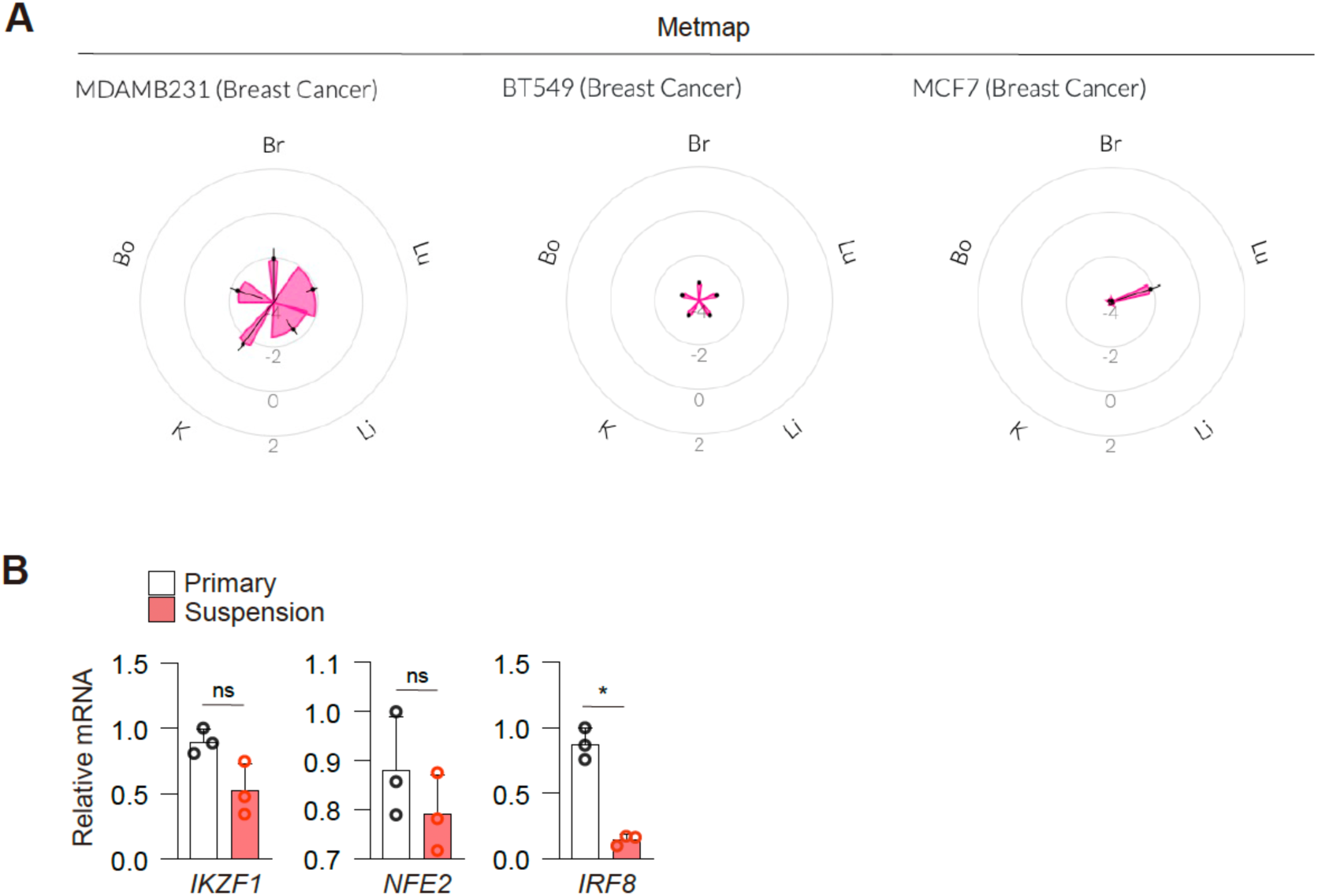
The ability to generate AST factors-expressing CTC-like cells is correlated with their metastatic capability. **A**, Petal plots showing the metastatic patterns of 3 breast cancer cell lines. Data for these plots were obtained from Jin, X., *et al*. (https://depmap.org/metmap). **B**, Quantitative real-time PCR analysis of AST factor transcript levels in cells harvested from suspension assay.

**Supplementary Figure 2.**
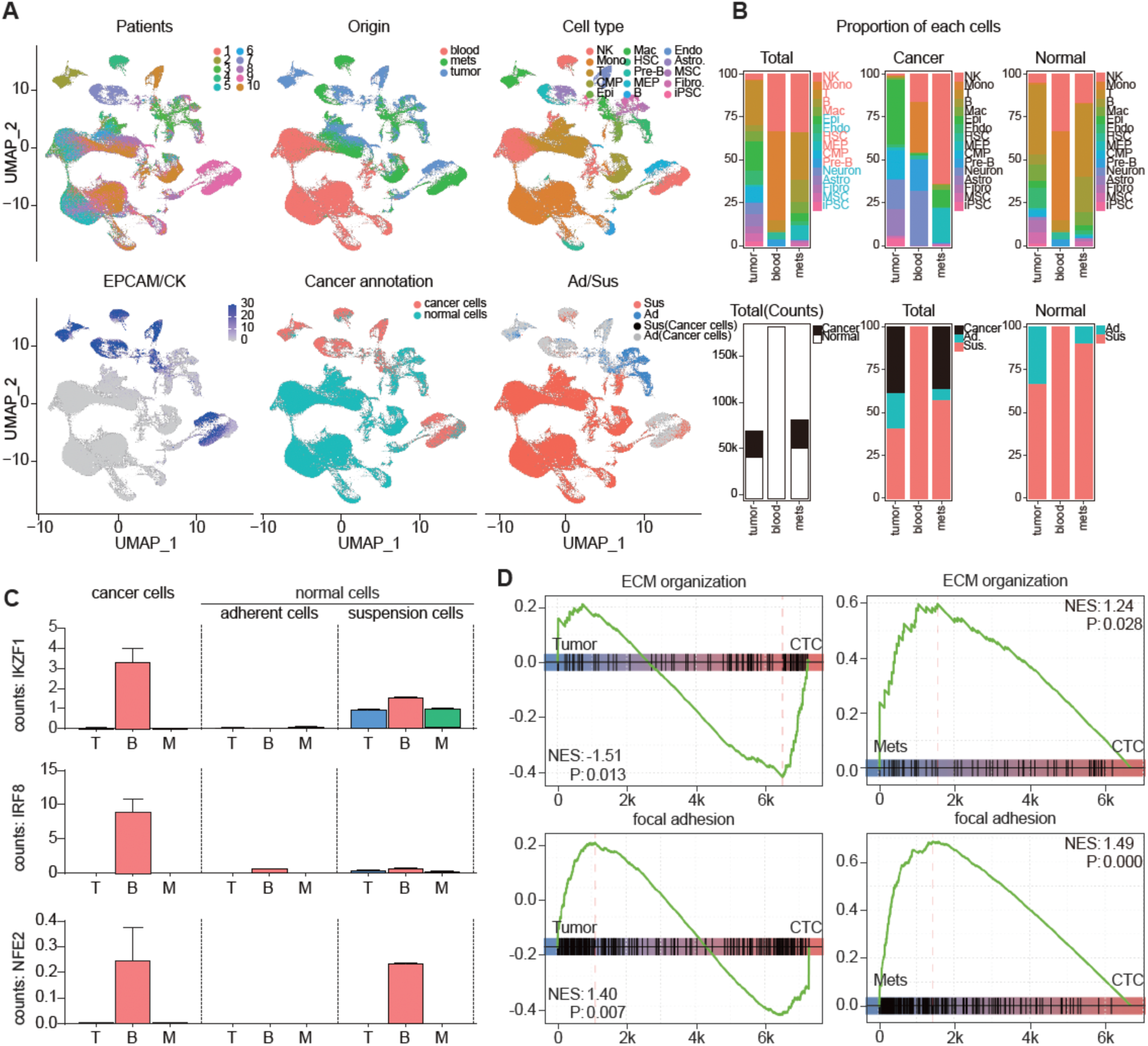
Comprehensive overview of single cell RNA sequencing data from breast cancer patients. **A**, Uniform manifold approximation and projection (UMAP) of scRNA-seq cells (n = 316,539 cells) colored by patient, origin, cell type, normalized gene expression of cancer cell markers (EPCAM, KRT5, KRT7, KRT8, KRT18, and KRT19), cancer/non-cancer annotation, and the combined cancer/non-cancer and adherent/suspension annotation. **B**, Bar plots showing a breakdown of cell type composition across tumor, blood, and metastasis samples. The first row shows the proportion of various cell types in total cells, cancer cells, and normal cells. The second row shows the proportion of cancer cells versus normal cells, adherent versus suspension cells within cancer and normal cells, and adherent versus suspension cells within normal cells. **C**, Dynamite plots depicting the expression of the AST factors IKZF1, IRF8, and NFE2 in cancer cells from primary tumor, blood, and metastases, as well as normal cells split into adherent and suspension states. **D**, Gene set enrichment analysis (GSEA) plots for focal adhesion and ECM organization gene sets comparing CTCs to primary tumor or metastatic cells.

**Supplementary Figure 3.**
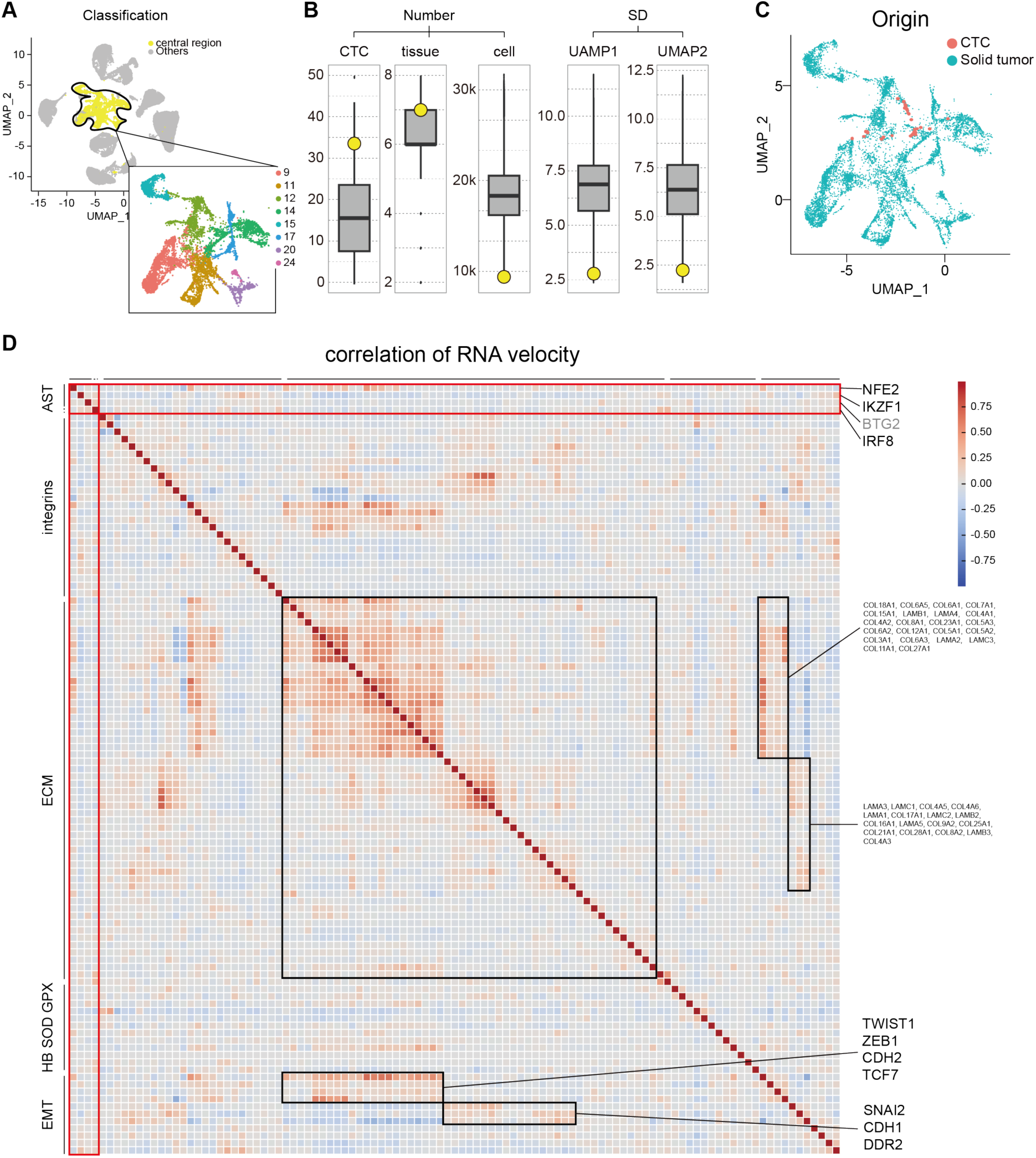
Central region-specific cell populations and CTC–solid tumor divergence in scRNA sequencing data. **A**, UMAP of cancer cells categorized into the central region and others. A magnified view of the central region is also shown to highlight its specific clusters. **B**, Boxplots of 1,081,575 cluster combinations (binomial coefficient with n = 25, k-permutation = 8) presenting numbers of CTCs, originating tissues, and total cells with standard deviations (SDs) for UMAP1 and UMAP2. Yellow dot, selected cluster combination; CTCs, n = 34, 11.93%; originating tissues, n = 7, 8.83%; total cells, n = 8,857, 99.31%; UMAP1 SD, 2.77, 90.69%; UMAP2 SD, 2.49, 0.36%. **C**, UMAP of cells in the central region colored by anchorage status. Turquoise, solid tumor; orange, CTCs. **D**, Heatmap of RNA velocity correlations among AST factor, ECM, integrin, superoxide resistance genes, and EMT genes.

**Table S1.** Overview of patient information used for single cell transcriptome analysis.

| Patient # | Metastasis tissue | Subtype | Total cell counts |  |  | Cancer cell counts |  |  |
| --- | --- | --- | --- | --- | --- | --- | --- | --- |
|  |  |  | Tumor | Blood | Meta | Tumor | Blood | Meta |
| patient1 | liver | Her2 | 1,746 | 7,209 | 2,411 | 659 | 2 | 1,033 |
| patient2 | neck | Luminal-B | 13,246 | 9,499 | - | 9,198 | - | - |
| patient3 | lung | Her2 | 14,254 | 25,180 | 11,982 | 516 | 1 | 2,531 |
| patient4 | lung | Luminal-B | - | 25,394 | 10,900 | - | 2 | 7,507 |
| patient5 | brain | Her2 | 1,569 | 19,071 | - | 82 | 1 | - |
| patient6 | lung | TNBC | 3,612 | 13,166 | 5,034 | 324 | 1 | 19 |
| patient7 | lymph | Luminal-A | 9,572 | 26,768 | 9,404 | 4,438 | 8 | 5,833 |
| patient8 | neck | TNBC | 11,741 | 9,117 | 12,746 | 1,257 | 7 | 13 |
| patient9 | liver | Luminal-A | 8,077 | 21,471 | 16,458 | 5,032 | 9 | 11,641 |
| patient10 | lymph | TNBC | 5,795 | 14,629 | 10,138 | 4,721 | 19 | 13 |
|  |  |  | 69,612 | 171,504 | 79,073 | 26,227 | 50 | 28,590 |

**Movie S1. Dissemination assay of various breast cancer cell lines**

**A**, Video of MCF7 cell proliferation in the dissemination assay.

**B**, Video of BT549 cell proliferation in the dissemination assay.

**C**, Video of HS578T cell proliferation in the dissemination assay.

**D**, Video of MDA-MB-231 cell proliferation in the dissemination assay.

